# Evaluating Graph Neural Network Architectures for Multi-Omics Cancer Subtyping using Methylation and Gene Expression Profiles

**DOI:** 10.64898/2026.08.21.745839

**Authors:** Julia Schirmacher, Miriam C. Maurer, Jacqueline M. Metsch, Hryhorii Chereda, David B. Blumenthal, Anne-Christin Hauschild

**Affiliations:** Department of Medical Informatics, University Medical Center Göttingen, Germany; Campus Institute Data Science (CIDAS), University of Göttingen, Germany; Biomedical Network Science Lab, Department Artificial Intelligence in Biomedical Engineering, Friedrich-Alexander-Universitä Erlangen-Nürnberg, Germany; Department of Predictive Deep Learning for Medicine and Healthcare, Justus-Liebig University Gießen, Germany

**Keywords:** Graph Neural Networks, Protein-Protein-Interaction Networks, Deep Learning, Graph Convolutional Network, Graph Attention Network

## Abstract

**Motivation:** Graph Neural Networks (GNNs) have gained increasing interest in the biomedical domain, as the integration of prior knowledge and deep neural networks has the potential to enhance insights into molecular processes and disease mechanisms. However, a comprehensive and systematic assessment of model architectures, data modalities, graph structures, and their performance for graph signal classification in the biomedical domain is yet to be performed. In order to close this gap, we conducted a benchmarking study on multiple GNNs on a Protein-Protein Interaction (PPI) network for Kidney Renal Clear Cell Carcinoma and Breast cancer subtype prediction, performing an in-depth investigation of architectures, incorporating skip connections and various data modalities.

**Results:** While none of the GNNs outperforms the structure-agnostic Multi-Layer Perceptron baseline, all of them can handle bimodal data (gene methylation and expression) and offer the ability to gain explainability based on PPIs. We offer practical guidelines for applying GNNs to graph signal processing tasks specifically for cancer classification. Depending on the underlying dataset and PPI structure employed, models on different data modalities outperform others. Overall, we suggest using ChebNet, which tends to outperform the Graph Convolutional Network and the Graph Attention Network in cancer subtype prediction. We recommend using GNN architectures that employ a simple flattening readout layer, as they provide better classification performance and faster training time than those with global average pooling. Additionally, we tested residual connections, but they had only an insignificant impact on classification performance.

**Availability and implementation:** Code and data available at https://github.com/HauschildLab/GNN4PPI.

**Supplementary information:** Supplementary data attached.

## 1. Introduction

High-throughput next-generation sequencing technologies are increasingly being used in clinical practice to aid in cancer prognosis. However, single-modality data analysis provides only a narrow view of cellular functions, limited to a single molecular level. Thus, systems biology, in contrast, aims to integrate multi-modal information across different Omics platforms, including, e.g., genomics, epigenomics, transcriptomics, proteomics, and metabolomics, providing the opportunity to understand causal relationships across multiple levels of cellular organization (Menyhárt and Győrffy, 2021). Moreover, additional information is available on structural relationships of such biological entities, such as gene regulatory networks, signal transduction graphs, sequence similarity networks, and Protein-Protein-Interaction (PPI) networks (Koutrouli et al., 2020). This biological knowledge and its underlying structure can be represented in various forms, including graphs, where nodes represent entities and edges define their relationships.

Recent studies have shown promising results using graph neural networks (GNNs) to integrate such biological insights in multiomics cancer research, including subtype prediction (Zohari and Chehreghani, 2025; Rhee et al., 2018). Additionally, this lays the foundation for future explainable AI (xAI) studies to enhance biological insights, like Chereda et al. (2024, 2021); Pfeifer et al. (2022); Metsch et al. (2024). However, the plenty of complex GNN architectures, such as Graph Convolutional Network (GCN) (Kipf and Welling, 2017), Graph Attention Network (GAT) (Veličković et al., 2018), and ChebNet (Defferrard et al., 2017), implementation strategies and parameters in combination with the underlying multi-omics datasets require a careful selection and optimization. Despite numerous general benchmarking studies on GNNs in other domains, there is a significant gap in comprehensive evaluations and comparisons of these models in systems medicine, particularly for specialized domains and clinical tasks. While basic GNN model comparisons exist, see Ramirez et al. (2020) and Alharbi et al. (2025), these often do not include a baseline model or lack a thorough investigation of crucial architectural GNN variations.

Therefore, this pilot study aims to fill this gap by (1) comparing GNN architectures in general with a standard multilayer perceptron (MLP), (2) evaluating the usability of different multi-omics datasets (gene expression, methylation, and full data), and finally (3) evaluating the potential of different GNN architectures (GCN, GAT, ChebNet), as well as the effect of the readout function and residual connections following the idea of Li et al. (2023) within each architecture. Additionally, this benchmark incorporates a standardized nested model selection, training convergence evaluation, detailed performance analysis per data split, and extensive test repetitions for solid statistical analysis.

## 2. Methods

### 2.1. Dataset

The benchmark evaluation was carried out on two cancer datasets. In both cases, a fixed graph topology is shared across all patients, while node features are defined by patient-specific data. This topology represents interactions among the features and incorporates prior biological knowledge in the form of a protein-protein interaction (PPI) network. Patient multi-omics profiles form multi-modal graph signals to train GNNs for the classification task.

The first dataset, the KIRC dataset, follows the setup described in Pfeifer et al. (2022) and Metsch et al. (2024). Specifically, this means that the structural basis is a PPI network obtained from the STRING database (Szklarczyk et al., 2021). Then, patient-specific node features for 306 Kidney Renal Clear Cell Carcinoma (KIRC) patients and 200 controls with random cancer types are incorporated. The multimodal features comprise mRNA gene expression and DNA methylation data obtained from The Cancer Genome Atlas (TCGA). The data are harmonized and filtered by cancer-relevant genes as proposed in Schulte-Sasse et al. (2021). Genes with missing values in any patient are removed, followed by *min-max* normalization. After reducing to its main connected component, the resulting PPI networks consist of 1 594 nodes with two-dimensional features and 2 560 edges.

The second dataset, the BRCA dataset, again consists of a universal PPI network structure, here taken from IID (Kotlyar et al., 2025), and patient-specific methylation and gene expression data from TCGA. This time, they originate from 360 luminal A breast cancer patients and 329 patients suffering from breast cancer type luminal B, basal-like, HER2-enriched, or normal-like. The gene expression data, which are normalized using the gene length corrected trimmed mean of M-values (GeTMM) method and are log_2_(*x* + 1) transformed, originate from a previous study (Chereda et al., 2024). The harmonized gene-wise methylation data (HM450K) are downloaded from linkedomics.org (Vasaikar et al., 2018). The patients present in both data sources were matched, and genes with at least one missing value were filtered out. Then, we conducted a *min-max* normalization. To focus on relevant genes only, the PPI network was filtered by breast cancer genes with interactions proven in at least 3 experiments based on IID filters. Finally, the remaining genes with omics and PPI network information are matched, and the main connected component of the resulting network is extracted. It consists of 1 917 nodes and 6 496 edges with two-dimensional patient-specific node features.

### 2.2. Experiments

In this study, a baseline model and three GNN architectures are evaluated on multiple data modalities and with different architectural settings using a PPI network as the structural foundation for cancer prediction. A structure-agnostic baseline model, i.e., a multi-layer perceptron (MLP), and three GNN models, Graph Convolutional Network (GCN) (Kipf and Welling, 2017), Graph Attention Network (GAT) (Veličković et al., 2018), and ChebNet (Defferrard et al., 2017) are analyzed. All models are tested independently using identical data splits and training procedures, ensuring the strict separation between model selection and evaluation.

The setup can be split up into 3 experiments.

#### Experiment 1

In the first experiment, a general comparison between GNNs and the baseline model, MLP, is drawn. For this, the models are evaluated across different settings, including different data modalities and readout functions, to gain an overview of the capability of GNNs compared to the MLP.

#### Experiment 2

Then, the effect of different data modalities on the model performances is analyzed in more detail. This means that the performance of the MLP and GNNs is evaluated using either the multi-omics feature nodes, including gene expression and methylation data, or only the corresponding single-omics data each.

#### Experiment 3

Finally, several aspects of GNN architectures are considered.

a. First, the performance differences among the GNN models, the GCN, GAT, and ChebNet, are analyzed.
b. Then, the effect of the readout function, the aggregation function from the resulting node features to the single vector representing the entire graph, is examined. Here, global average pooling or simple flattening is considered.
c. In the end, the impact of integrating residual connections is investigated. This includes comparing MLPs and GNNs in their plain form and with residual or dense connections added. This is only conducted on the KIRC dataset, headed by the results on these data and the comparatively high cost of this sub-experiment.

The experimental setup is visualized in Figure 1.

**Fig. 1:**
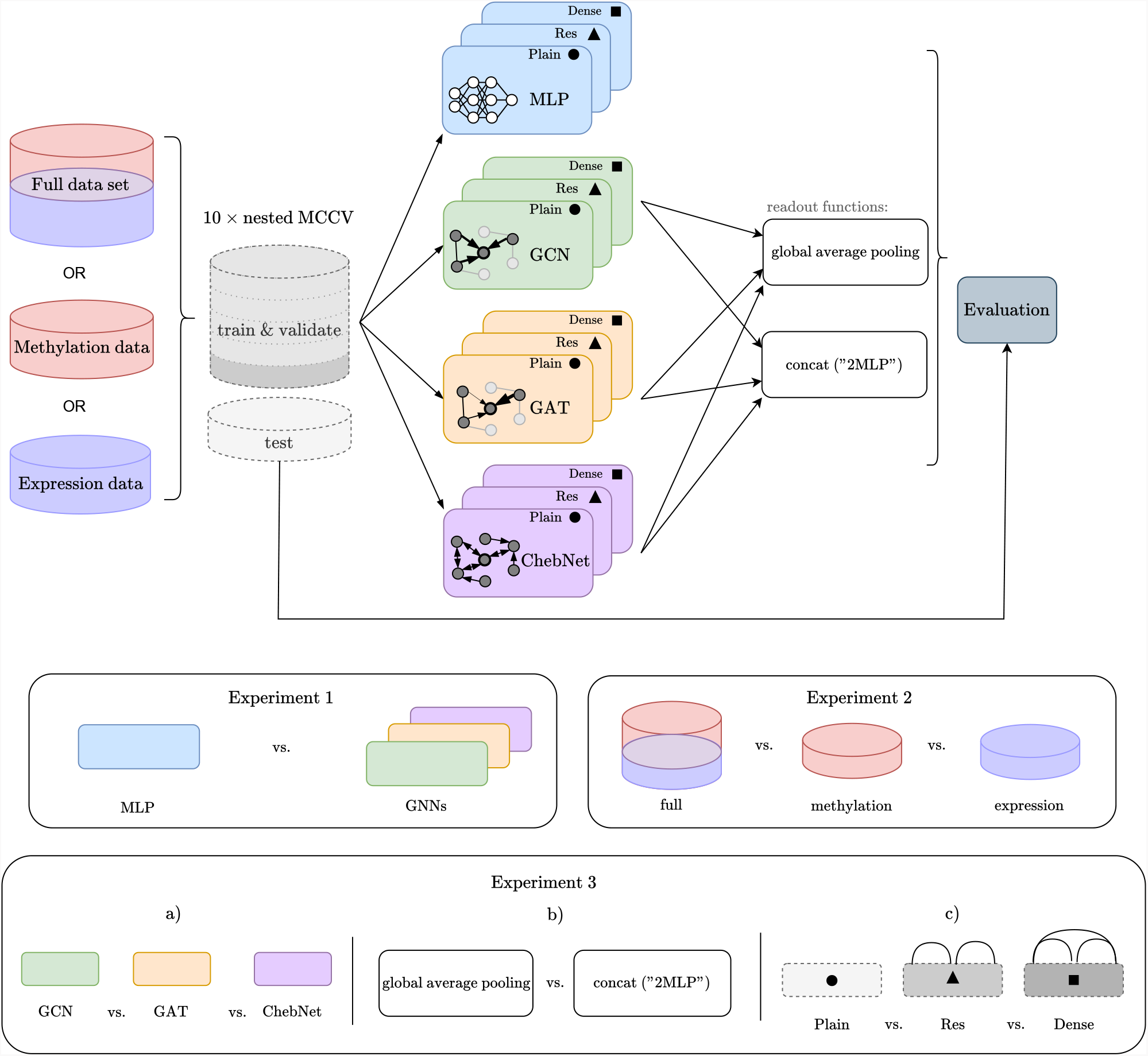
The general workflow of the benchmark. Different data modality and model architecture combinations are evaluated and analyzed in 3 experiments. Generally, different data modalities (multi-omics or single omics) are employed. One baseline MLP and three GNNs, namely a GCN, GAT, and ChebNet, are evaluated. All models can be run in their plain version or with residual or dense connections added. As a readout function, either global average pooling or simple flattening of the node features is used. After training and validation, all models are evaluated on identical test data splits. Data are split ten times using nested Monte-Carlo cross-validation; details are provided in Section 2.4 and visualized in the Supplementary Figure 1. In experiment 1, the performance of the MLP is compared with the GNNs’ performance generally. Then, in experiment 2, the impact of the underlying data modality is studied. In experiment 3, details of the model architectures are compared, namely a) the general model, b) the readout function, and c) the incorporation of residual or dense connections.

### 2.3. Models

#### Baseline model MLP

The used baseline model is a simple MLP that operates on the concatenated node features. It is not permutation-invariant, however, since the graph structure is fixed, the node features can be consistently ordered, thereby allowing the model to process graph data. The architecture of the MLP is comparable to that of the GNNs, which both have 2 hidden layers, 64 hidden features, and ReLU activation.

#### GCN

The GCN (Kipf and Welling, 2017) is a GNN derived from spectral graph theory, with constraints and simplifications applied to graph convolutions formulated as filtering operations in the Fourier domain. While single filter operations are simplified, complex filtering is achieved by stacking multiple filters in layers within a neural network. The operation can also be interpreted spatially as an aggregation of neighboring node features on the graph structure. Following the original architecture by Kipf and Welling (2017) and findings from Shchur et al. (2019) and Kim and Cho (2021), the GCN has 2 hidden layers with 64 hidden features each and ReLU activation.

#### GAT

The GAT (Veličković et al., 2018) represents an extension to the GCN approach by incorporating a self-attention mechanism. This allows the network to adaptively weight neighboring features during the aggregation process. This is achieved with a single-layer feedforward network, LeakyReLU, and softmax for normalization. In accordance with the original architecture in Veličković et al. (2018) and the findings from Shchur et al. (2019), the GAT architecture consists of 2 layers with 8 attention heads, each having 8 hidden features and ELU activation.

#### ChebNet

The ChebNet (Defferrard et al., 2017) builds on the spectral graph theory similar to the GCN, but with fewer simplifications and thus is a more general form of the GCN. Concretely, this means that a single filtering operation is simplified to a localized polynomial filter parametrized with Chebyshev polynomials for fast filtering. The resulting *K*-localized filter can be interpreted in the spatial domain as an aggregation of the node features in the *K*-hop neighbourhood. Additionally, the original architecture in Defferrard et al. (2017) also includes graph coarsening and pooling similar to downsampling in CNNs, which is, for the sake of comparability of the filter itself, not considered in the following, but will be evaluated in the future. Similar to the architectures in Defferrard et al. (2017) and Chereda et al. (2024) and in accordance with the previous models, 2 hidden layers with 64 hidden features, *K* = 8, and a ReLU activation were employed.

#### ResNet and DenseNet

To address gradient vanishing and potentially over-smoothing in GNNs (Zhou et al., 2020), the idea of ResNet (He et al., 2016) and DenseNet (Huang et al., 2017) is integrated in this benchmark. ResNet employs skip connections, whereby hidden features are transferred to a following layer via addition. Conversely, DenseNet uses a concatenation of hidden feature vectors with those of all subsequent layers. While the efficacy of these techniques has been demonstrated for GNNs (Li et al., 2023), further examination is necessary for the specific task and setting under consideration. In the Dense variant, the feature size is reduced by half from the second layer onwards in order to mitigate the rapid dimensional growth due to concatenation. In both variants, one layer is skipped. To ensure comparable model sizes and following findings of Li et al. (2023), architectures with 3 layers and 64 hidden features are employed for all variants since not all variant differences are evident with fewer layers. Other structural choices remain unaltered by these variants when combined with different model types.

#### Readout

The GNNs described above output node representations, but to obtain a graph embedding that can be used as input for a final classifier, a transformation, the so-called readout, is needed. One widely used standard approach is the dimension-wise mean of the features, which results in a vector with a dimensionality equal to the *hidden feature size*; this is also called global average pooling (Wu et al., 2021). Another approach, which is not permutation-invariant but can be applied here since we are examining graph signal classification (i.e., the number of nodes is fixed, and thus the nodes can be ordered), is the simple flattening of the features, meaning a concatenation of the node features (Buterez et al., 2022). This readout is denoted as “2MLP” in the following to emphasize the direct transfer to the classifier layer. Both readout options are followed by a single feedforward classifier layer with an input width of either the *hidden feature size* or #*nodes × hidden feature size*, respectively, and a single output unit for binary classification. This difference in the final layer’s input width is the primary reason for the huge difference in the number of parameters between the global average and “2MLP” model variants.

### 2.4. Evaluation

To ensure fair benchmarking, a consistent training setup is utilized across all models, including Binary Cross Entropy loss, the Adam optimizer (Kingma and Ba, 2017), and Glorot initialization (Glorot and Bengio, 2010). Regularization techniques, including dropout, weight decay, and early stopping, are implemented. The former two and the learning rate are realized as hyperparameters. The GAT model introduces an additional hyperparameter, a dropout rate integrated into the self-attention mechanism. Hyperparameter choices were reduced from 5 to 3 per model in a pretest. See Supplementary Table 3 and Table 4 for the ranges used in the final evaluation. Early stopping is implemented with a patience of 100 epochs, and the maximum number of epochs is 10 000, which is rarely even reached, see Supplementary Figure 3 and Figure 5, but facilitates flexible training. To ensure the separation of model selection and assessment and to provide separate validation data sets for early stopping, a nested Monte-Carlo cross-validation scheme was implemented. It consists of 10 independent, random outer data splits in an outer test set (20%) and an outer training/validation set (80%). The outer training/validation set is further split using 5-fold cross-validation for early stopping, resulting in 50 repetitions for model assessment. Additionally, the outer training/validation set is split beforehand for model selection, which aims to choose the best hyperparameter configuration based on the highest AUC. Therefore, another 5-fold cross-validation is performed with an inner test set and an inner training/validation set, randomly split into an inner training set (80%) and a validation set (20%) for early stopping. This results for MLP, GCN and ChebNet in 5 · 3^3^ = 135 and for GAT, due to an additional hyperparameter, even 5 · 3^4^ = 405 inner model training repetitions per test set. This procedure is visualized in the Supplementary Figure 1. Additionally, Supplementary Table 5 shows the number of parameters for the models in all different settings exemplarily for the KIRC data. The experiments are run on an HPC server with 4*×*Nvidia A100 (VRAM 40 GiB, 2*×*CPU Zen3 EPYC 7513, RAM 12 GiB) per node in parallel, one GPU per training run each. The benchmark implementation is available at https://github.com/HauschildLab/GNN4PPI.

## 3. Results

This section presents the key results of the three experiments on the two datasets and provides additional insights into practical aspects, such as model training time.

### 3.1. Experiment 1

The results of experiment 1, the overall performance comparison of the MLP and the GNNs, are shown in Figure 2(a) and Figure 3(a). It illustrates the performance of the plain MLP and GNN models across three data modalities in terms of Matthew’s correlation coefficient (MCC). It reveals that GNNs generally achieve performance similar to that of MLPs. This is accomplished for all models, even if some GNN models lag behind on the single-omics data modalities.

**Fig. 2:**
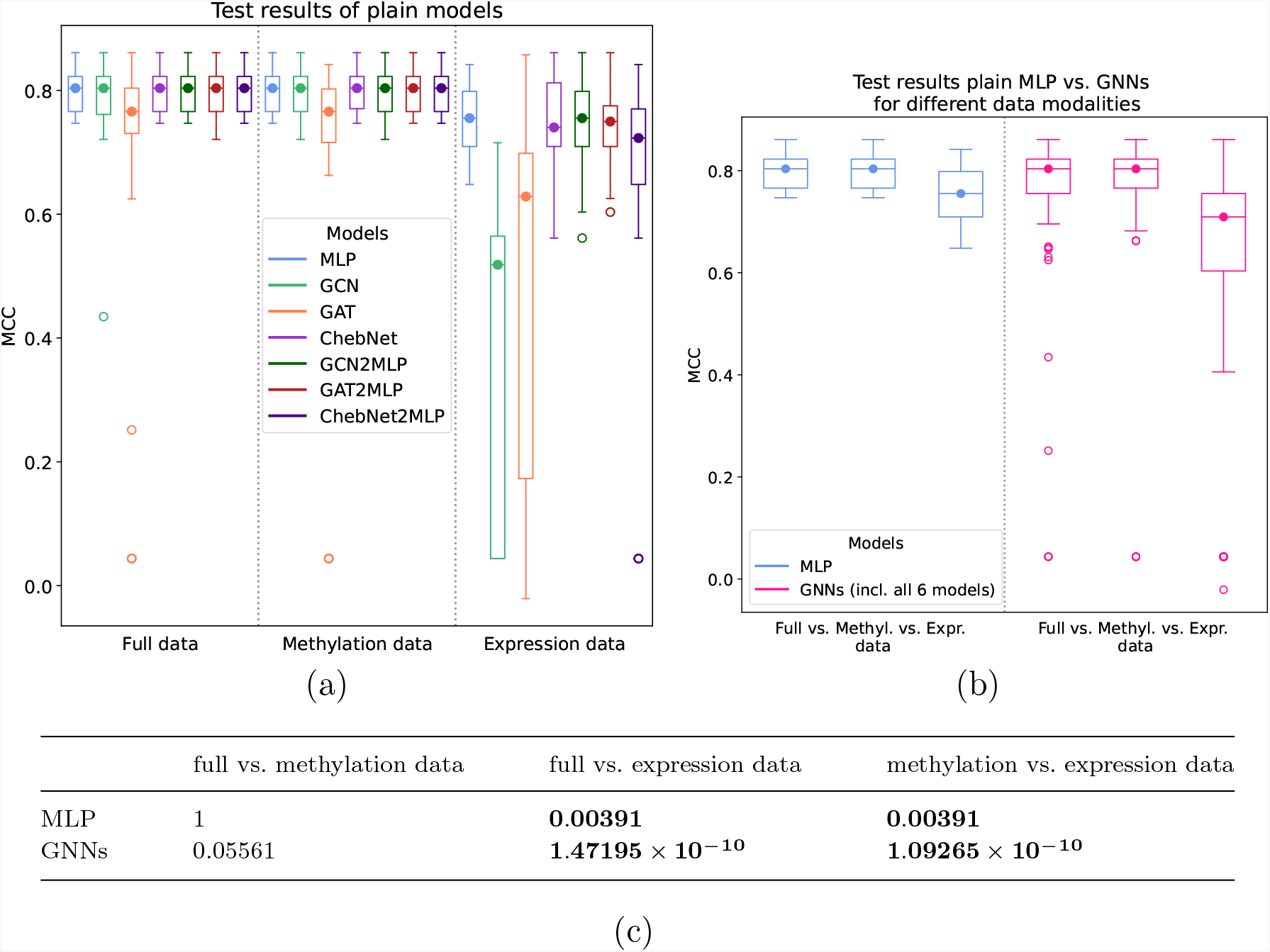
Results on the **KIRC data:** The MCC results distribution and p-value of the Wilcoxon signed rank test (on the mean test performance per test set) of the models aggregated regarding the experiment question 1 and 2. Here, the results of all 50 outer runs (10 test sets with 5-fold cross-validation each) are depicted as box plots. P-values under the significance level *α* = 0.01 are marked in bold. (a) Experiment 1: Overview of all plain models for different data modalities. (b), (c) Experiment 2: Comparison between models on different data modalities, the full data, or the methylation or expression data; here, all 6 different GNN models are combined.

**Fig. 3:**
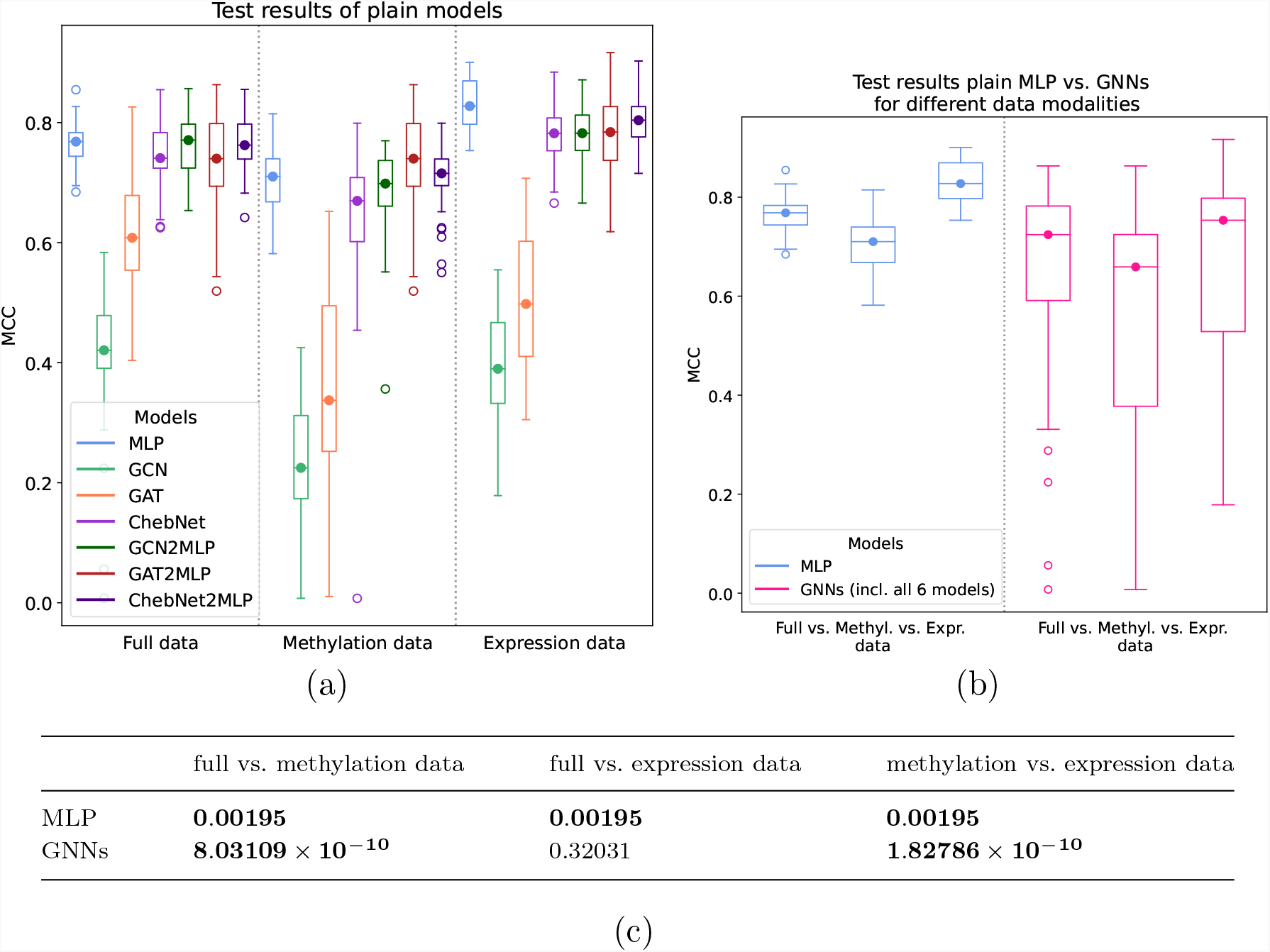
Results on the **BRCA data:** The MCC results distribution and p-value of the Wilcoxon signed rank test (on the mean test performance per test set) of the models aggregated regarding the experiment question 1 and 2. Here, the results of all 50 outer runs (10 test sets with 5-fold cross-validation each) are depicted as box plots. P-values under the significance level *α* = 0.01 are marked in bold. (a) Experiment 1: Overview of all plain models for different data modalities. (b), (c) Experiment 2: Comparison between models on different data modalities, the full data, or the methylation or expression data; here, all 6 different GNN models are combined.

### 3.2. Experiment 2

This trend can also be seen in the evaluation of experiment 2 in Figure 2(b), Figure 2(c), Figure 3(b), and Figure 3(c). The Figures show the MCC performance of the models, but this time all GNN models are gathered for a better overview of the performance across different data modalities. Additionally, the tables show the p-values for the Wilcoxon signed-rank test of the MCC test results, averaged across the outer folds, interpreted with a significance level of *α* = 0.01 to detect statistically significant performance differences.

KIRC data shows that the performance declines if only expression data are used for model training and prediction. Both the MLP and GNNs perform significantly better with the expression data or the full data underlying compared to the methylation data only. This trend is even more severe for GNNs, which we already ascertained in experiment 1.

In the breast cancer dataset, models performed best when trained exclusively on expression data, while incorporating the full dataset resulted in a decrease in performance. Models trained solely on methylation data performed the worst. This contrasts with the findings reported by the KIRC data, where performance declined when only expression data were used.

### 3.3. Experiment 3

In contrast, experiment 3 evaluates the performance differences of different architectures. Figure 4(a), Figure 4(d), Figure 5(a), and Figure 5(c) show the results of experiment 3a) comparing the MCC results of GCN, GAT, and ChebNet over all data modalities. It reveals the superiority of the ChebNet compared to the GCN and GAT, which is also demonstrated to be statistically significant for both datasets. It stands out that the ChebNet not only has a higher median MCC, but also has fewer outliers than GCN and GAT. In experiment 3b), the readouts, i.e., the global average pooling and the feature flattening (“2MLP”), are analyzed. Figure 4(b), Figure 4(e), Figure 5(b), and Figure 5(d) show the results analogously to the previous. All of them show that for the GCN and GAT, the flattening model performs statistically significantly better and clearly with fewer severe outliers, whereas for the ChebNet, no huge differences are observed. In retrospect, Figure 2(a) show that for the KIRC data, this trend is particularly pronounced in the single-omics expression data modality. Finally, experiment 3c), which was only conducted on the KIRC data, presented in Figure 4(c) and Figure 4(f), suggests that residual and dense connections can not improve the performance for either the MLP or the GNNs.

**Fig. 4:**
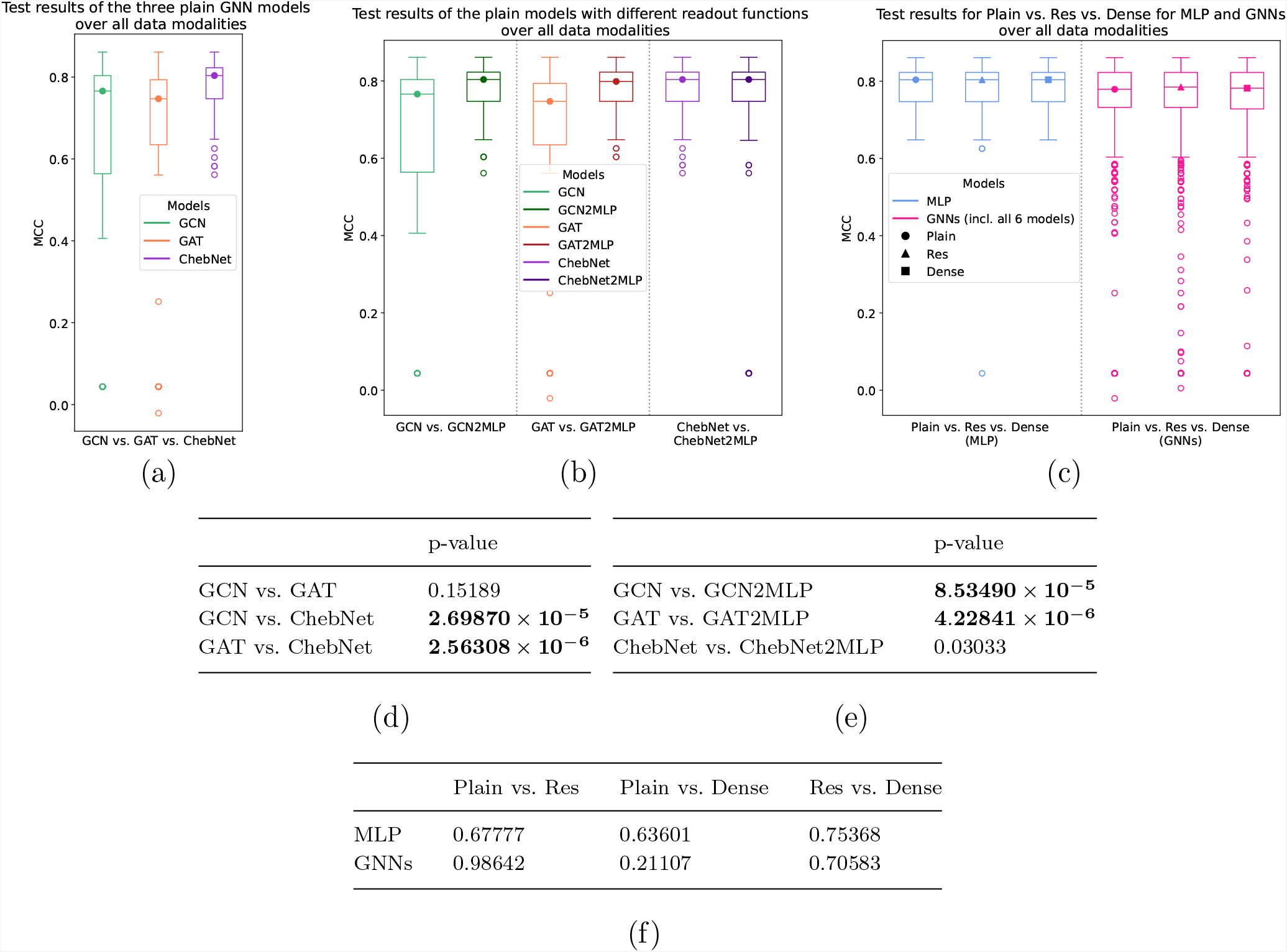
Results on the **KIRC data:** The MCC results distribution and p-value of the Wilcoxon signed rank test (on the mean test performance per test set) of the models aggregated regarding the experiment question 1 and 2. Here, the results of all 50 outer runs (10 test sets with 5-fold cross-validation each) are depicted as box plots. P-values under the significance level *α* = 0.01 are marked in bold. (a),(d) Experiment 3a: Comparison of the models GCN, GAT, and ChebNet; here, the model results on the different data modalities are combined. (b), (e) Experiment 3b: Comparison of the readout functions, i.e., global average pooling and flattening (called “2MLP”); here, the model results on the different data modalities are combined. (c), (f) Experiment 3c: Comparison between models without skip connections (Plain), with residual connections (Res), and with dense connections (Dense); here, all 6 different GNN models and the model results on the different data modalities are combined.

**Fig. 5:**
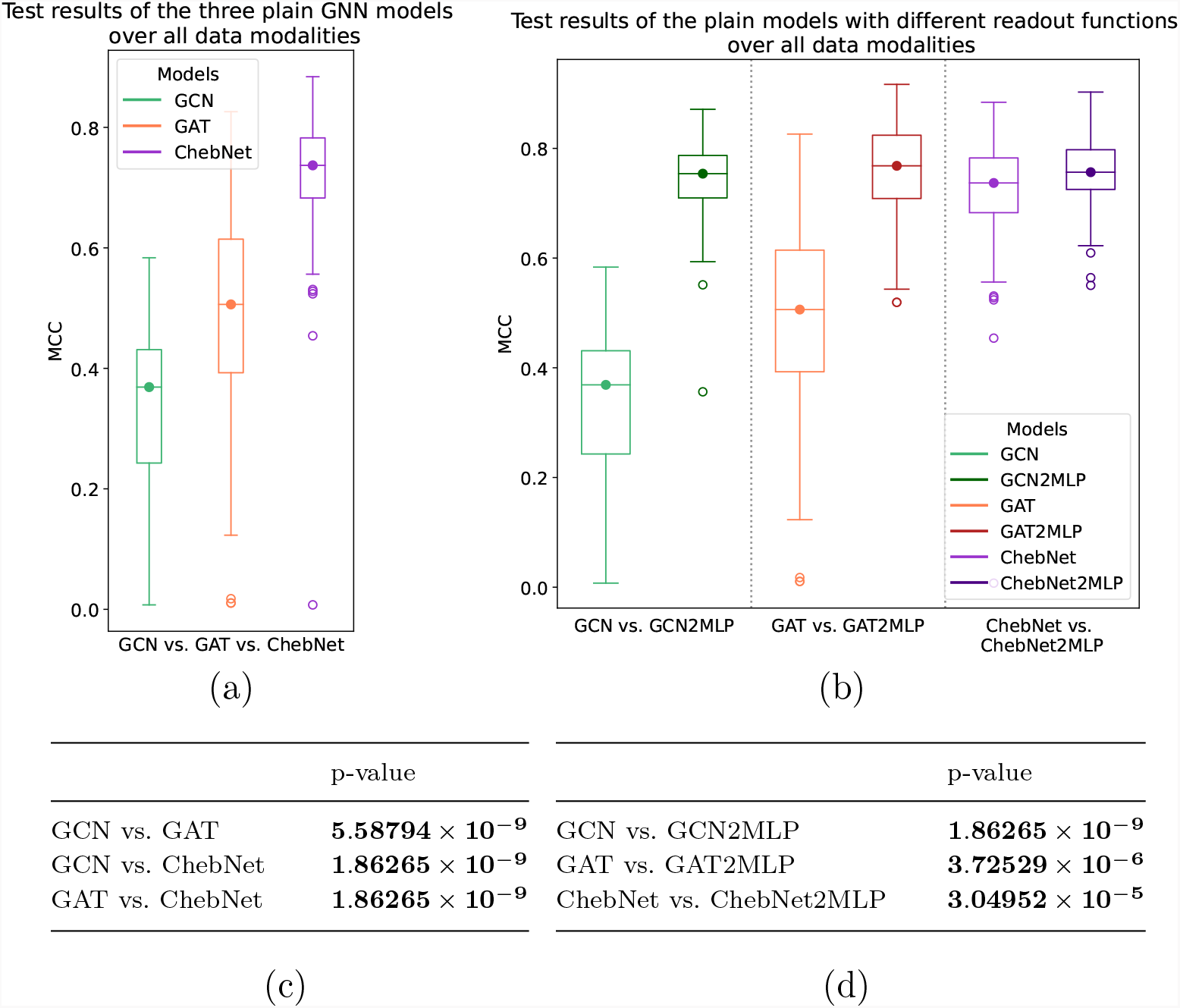
Results on the **BRCA data:** The MCC results distribution and p-value of the Wilcoxon signed rank test (on the mean test performance per test set) of the models aggregated regarding the experiment question 1 and 2. Here, the results of all 50 outer runs (10 test sets with 5-fold cross-validation each) are depicted as box plots. P-values under the significance level *α* = 0.01 are marked in bold. (a),(d) Experiment 3a: Comparison of the models GCN, GAT, and ChebNet; here, the model results on the different data modalities are combined. (b), (e) Experiment 3b: Comparison of the readout functions, i.e., global average pooling and flattening (called “2MLP”); here, the model results on the different data modalities are combined.

### 3.4. Practical aspects

Coming to the practical aspects, the model training time shown in Figure 6 and 7 reveals severe differences among the models and the underlying data modalities. Overall, the GNN models combined with the flattening readout are trained much faster than with the global average pooling readout. Moreover, the models trained on expression data seem to take longer to train than their corresponding methylation or full data models. In addition, the number of training epochs needed by the models, shown in Supplementary Figure 3 and Figure 5, reveals a similar trend for the number of epochs used. Again, for practical implications, the hyperparameters selected in the model selection procedure are depicted in Supplementary Figure 7 and Figure 9, and the performance on the wider hyperparameter grid during the pretest is shown in Figure 8 and Figure 10. Both suggest that the GNN models with global average pooling, especially the GCN and ChebNet, are more sensitive to the hyperparameter choice, as a smaller part of the hyperparameters in the grid result in high performance, and specific hyperparameter combinations are repeatedly selected in the inner model selection.

**Fig. 6:**
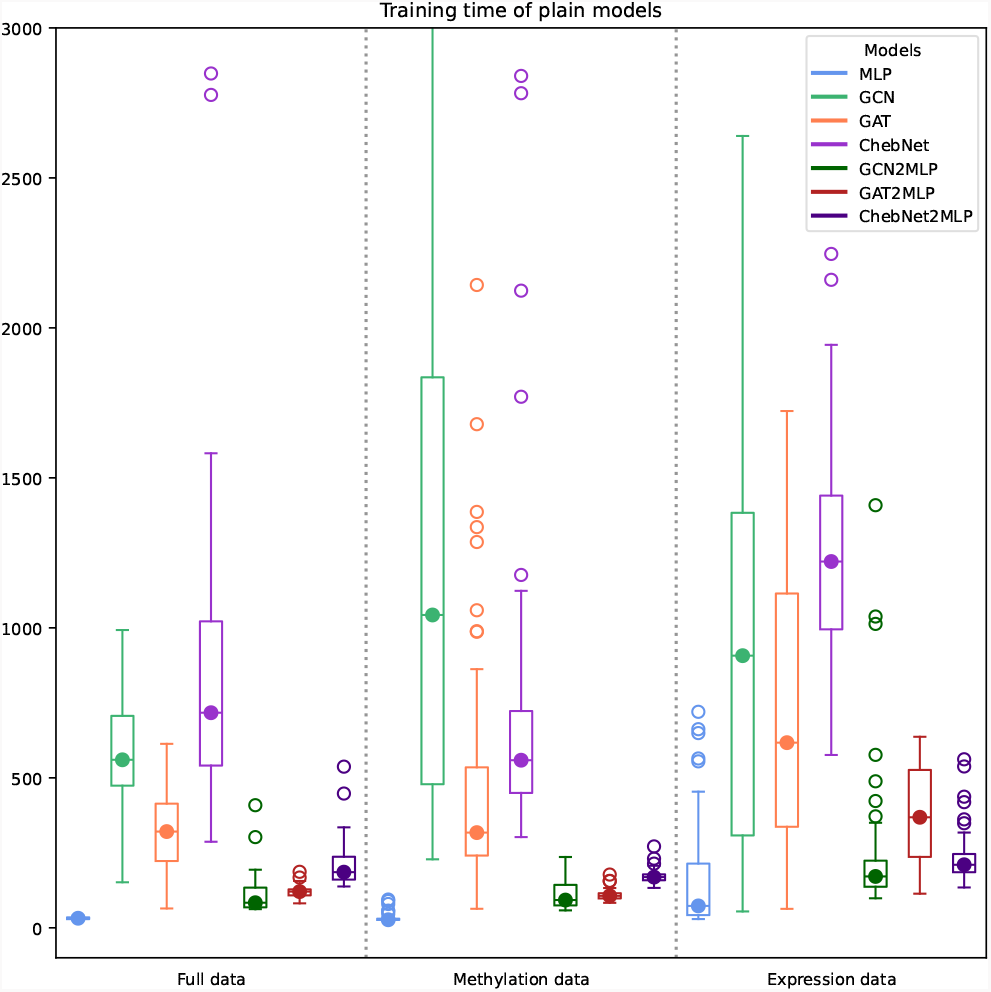
Results on the **KIRC data:** The distribution of training runtime per model training utilized per model architecture and data modality. For the used architecture, see Section 2.4.

**Fig. 7:**
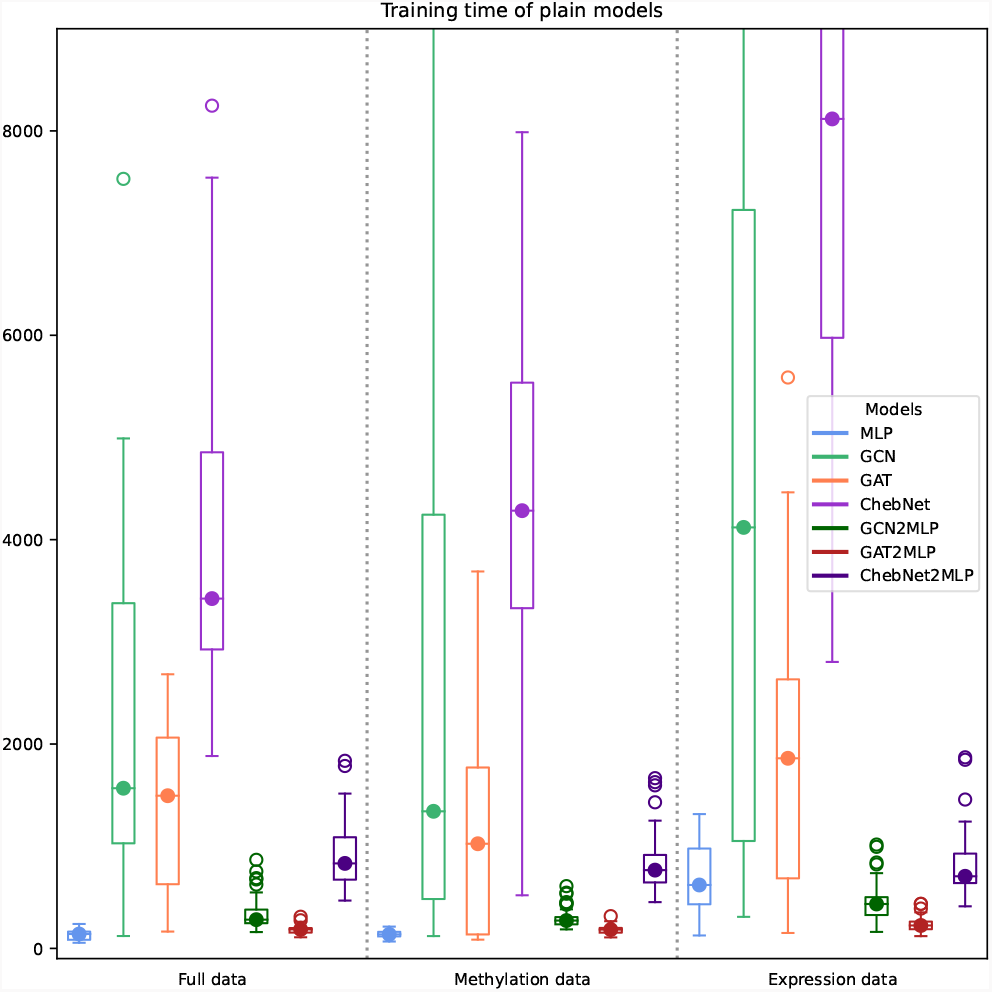
Results on the **BRCA data:** The distribution of training runtime per model training utilized per model architecture and data modality. For the used architecture, see Section 2.4.

## 4. Conclusion and Discussion

We have investigated the performance of GNNs and an MLP baseline for cancer classification based on four key factors: data modality, GNN type, readout strategy, and skip connections.

GNNs did not outperform the MLP baseline, despite the integration of PPI networks as prior knowledge to structure patient data, but still performed similarly. This finding leads to challenging the utility of both GNNs for cancer classification and PPIs as structural priors. While for the purely predictive tasks, GNNs utilizing PPI networks don’t seem to provide any leverage, a prior study (Chereda et al., 2024) established that ChebNet utilizing a PPI identifies more biologically significant features than standard MLPs. Thus, GNNs leveraging PPI networks are primarily advantageous when model predictions require biological explanation, which is an imperative in a clinical setting. Additionally, in this benchmark the number of model parameters are substantially lower for the GNNs compared to the MLP due to the huge input feature size for the MLP, which emphasizes, in view of the overall similar performance, the parameter efficacy of GNNs.

The comparison of the models based on different data modalities used for the training and prediction revealed that the superiority of models on a specific data modality differs between the datasets and PPI structures employed. For the task of distinguishing kidney cancer patients from those with other cancer types, the methylation modality is sufficiently informative, while the gene expression modality does not appear to improve classification performance further. This is in line with previous research (Pfeifer et al., 2022), where the authors explain that KIRC and other cancer types are similar at mRNA gene expression levels but differ primarily due to epigenetic factors. Moreover, Alharbi et al. (2025) found comparable results with a STRING PPI utilized and in a similar setting, but for multi-class classification. In contrast to the results from the KIRC dataset, models trained on the BRCA expression modality and its corresponding PPI network yielded superior results. This could be caused by the intrinsic selection bias resulting from the IID interaction association that is based on disease-specific transcription of genes in the database.

Still, Supplementary Figure 3 and Figure 5 demonstrate that multi-omics models tend to converge at a faster rate than single-omics models and can generally be trained faster, see Figure 6 and Figure 7. Therefore, while an additional modality may not improve classifier performance, its integration stabilizes training by improving noise handling and prediction verification.

Our results show that the GCN and GAT perform similarly for the given task, which was also found in Alharbi et al. (2025) for multi-class classification, but the ChebNet surpasses both. This can be justified by the fact that the ChebNet is a more general form of the GCN, but with more flexibility to adapt in training due to more parameters (Defferrard et al., 2017). Still, this is perceivable in the (for most variants) increased training time of the ChebNet.

We found that a simple flattening readout layer yields better classification performance for most of the GNNs compared to global average pooling. Global pooling leads to information loss because the information flow from all network nodes (representation vectors) collapses into a single node (single vector) without taking into account the graph structure. Our task at hand is classification of methylation or gene expression profiles of cancer patients structured by a PPI network. Since the PPI topology is constant across all patients, these profiles act as patient-specific graph signals. Consequently, our task is graph signal classification. In this scenario, a flattening readout is advantageous because the fixed number of nodes allows for a direct mapping of node representations. For a more generalized graph classification task, where the number of nodes can differ from one data point to another, a progressive graph coarsening with graph pooling is necessary to preserve graph structure information when computing its final representation (Duval and Malliaros, 2022). Thus, testing different graph pooling strategies for graph signal classification is our future work.

Finally, the model variants, including residual and dense connections, were demonstrated not to surpass the plain variants. However, Supplementary Figure 3 suggests that skip connections may facilitate convergence. This effect may also result from enhanced efficiency in training, but also increased model capacity in Res and Dense models due to the inclusion of an additional layer, and did not improve the performance of the models. This was also approved in the original study proposing GCNs (Kipf and Welling, 2017) with residual connections for node classification tasks. Still, for deeper GNNs, residual and dense connections may have the potential to increase model performance (Li et al., 2023). This remains to be investigated in the future. In addition, this missing performance improvement for GNNs with residual or dense connections was also shown for other data domains (Maurer et al., 2025).

Our benchmark focuses on the fair comparison of GNNs and a baseline MLP for cancer classification on gene expression and methylation data for the classification of KIRC and BRCA. This includes an extensive data splitting scheme. Retrospectively, as we can see in Supplementary Figure 4 and Figure 6, several inner and outer splitting repetitions were essential as the test sets seem to contain different difficulty levels, and also different early stopping sets can lead to fluctuating performance.

We omit benchmarking how underlying molecular network topologies affect GNN classification performance, as Alachram et al. (2021) reported that their effect on performance is negligible. However, while network topology may not significantly influence GNN performance metrics, it can alter patient-specific explanations (Chereda, 2022). The investigation of how explanations depend on network topology remains outside the scope of this study and is deferred to future work.

However, Ramirez et al. (2020) investigated multi-class cancer classification with ChebNet, including graph coarsening on a STRING PPI with or without added singleton nodes, meaning unconnected genes that are not included in the PPI but still have omics feature available. The addition of these singleton nodes, which can also include non-coding genes, resulted in a prediction accuracy increase of *>* 5%. This performance improve do not only emphasize the potential of GNNs on PPIs but also the weak point of PPIs of not capturing all gene regulations and activities (Ramirez et al., 2020). Incorporating this missing information needs to be further studied in the future.

Finally, we can derive practical recommendations for applying GNNs in the field of cancer classification with omics data and a fixed graph structure. First, for this graph signal classification task using methylation data and, if possible, expression data, is suggested. ChebNet was demonstrated to perform best and most stable and as a readout function, the so-called “2MLP”, i.e., the flattening of node features, is advised instead of global average pooling. This is not only recommended based on improved performance, but also for the seek of training time.

Combined with one of the numerous explainability methods for GNNs, such as GNNExplainer (Ying et al., 2019), PGMExplainer (Vu and Thai, 2020), or GraphLIME (Huang et al., 2020), or ChebNet-adapted LRP (Chereda et al., 2021), GNNs can not only provide solid model performance but also expand our knowledge and understanding of diseases of interests based on the underlying protein-protein interactions. This should be further investigated in the future, but this benchmark already represents a solid basis for model variant and data modality selection. Additionally, different types of PPIs and more data sources and cancer types will be investigated.

## 5. Competing interests

No competing interest is declared.

## 6. Author contributions statement

J.S., M.M., J.M., and A.H. conceptualized the project. J.S. performed all programming tasks as well as the conduction of experiments and analysis tasks. All authors performed review and editing.

## 7. Acknowledgments

The results shown here are in part based upon data generated by the TCGA Research Network: https://www.cancer.gov/tcga. This work is supported in part by funds from the German Ministry of Education and Research (BMBF) under grant agreements No. 01KD2208A and No. 01KD2414A (project FAIrPaCT). We gratefully acknowledge the computing time granted by the Resource Allocation Board and provided on the supercomputer Emmy/Grete at NHR@Göttingen as part of the NHR infrastructure, under the project *nib00044* and *nim00014*. In part funded by the Deutsche Forschungsgemeinschaft (DFG, German Research Foundation) – 405797229. In part this work is funded by the Bavarian State Ministry of Science and the Arts in the framework of the bidt Graduate Center for Postdocs.

## Notes

### Competing Interest Statement

The authors have declared no competing interest.

