## Supplemental Data for "Evaluating Graph Neural Network Architectures for Multi-Omics Cancer Subtyping using Methylation and Gene Expression Profiles"

#### Supplementary information

August 12, 2026

#### 1 Data splitting scheme

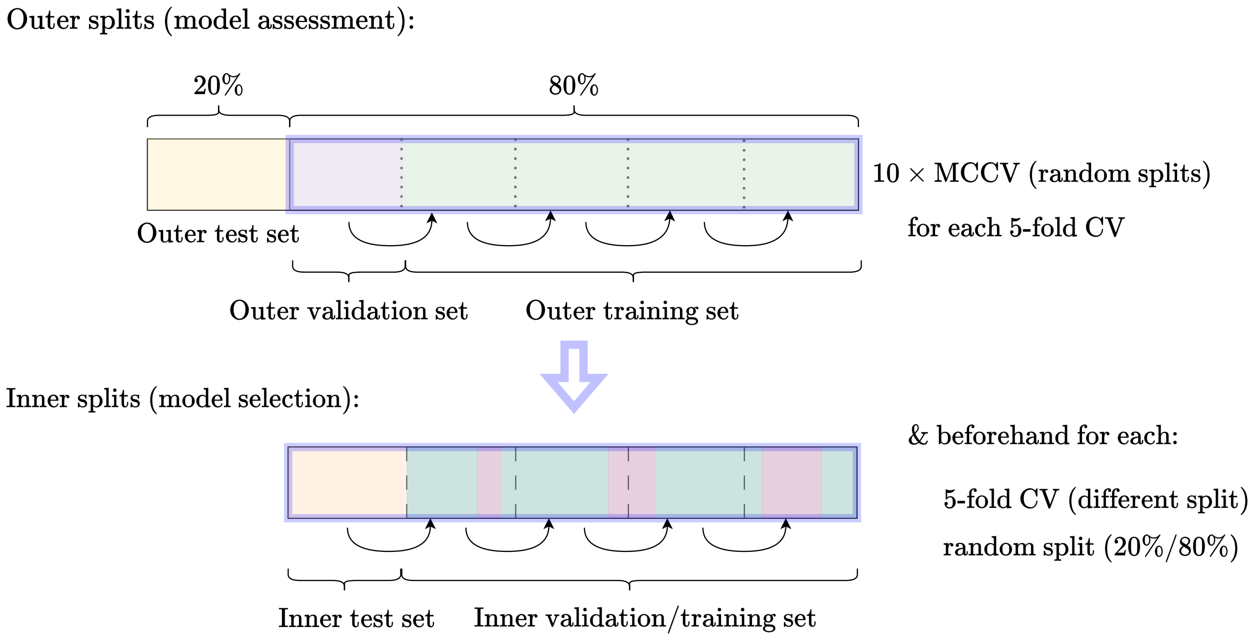


Figure 1: Data splitting scheme. The outer splitting refers to model assessment, the inner splitting to model selection. Per outer random test split, ten in number, a 5-fold CV with a validation set for early stopping is performed. Beforehand, in inner splitting a 5-fold CV is performed where the train/validation set is randomly split up again once.

### 2 Detailed performance results

#### 2.1 KIRC data

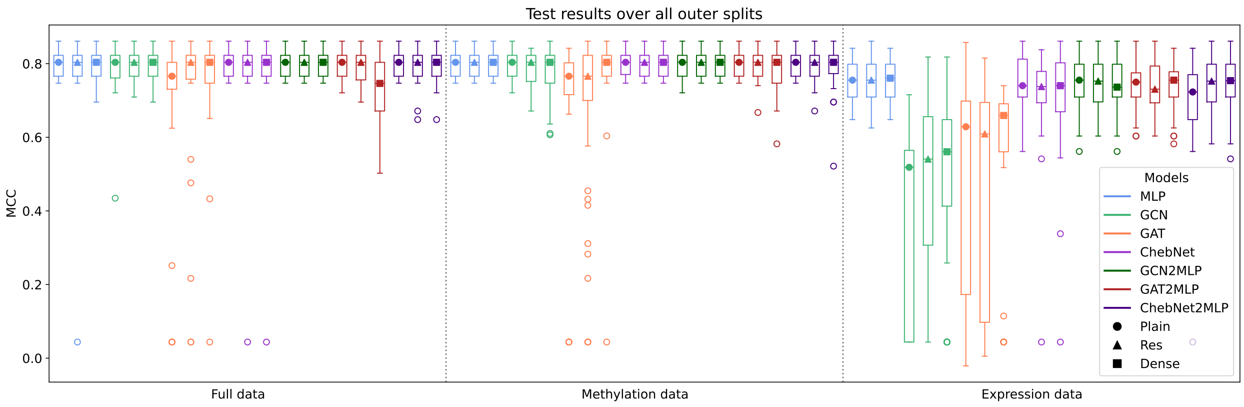


Figure 2: Experiment 3 for the **KIRC data**: The MCC result distribution of all models on all data sets. The results of all 50 outer runs (10 test sets with 5-fold cross-validation each) are depicted as box plots.

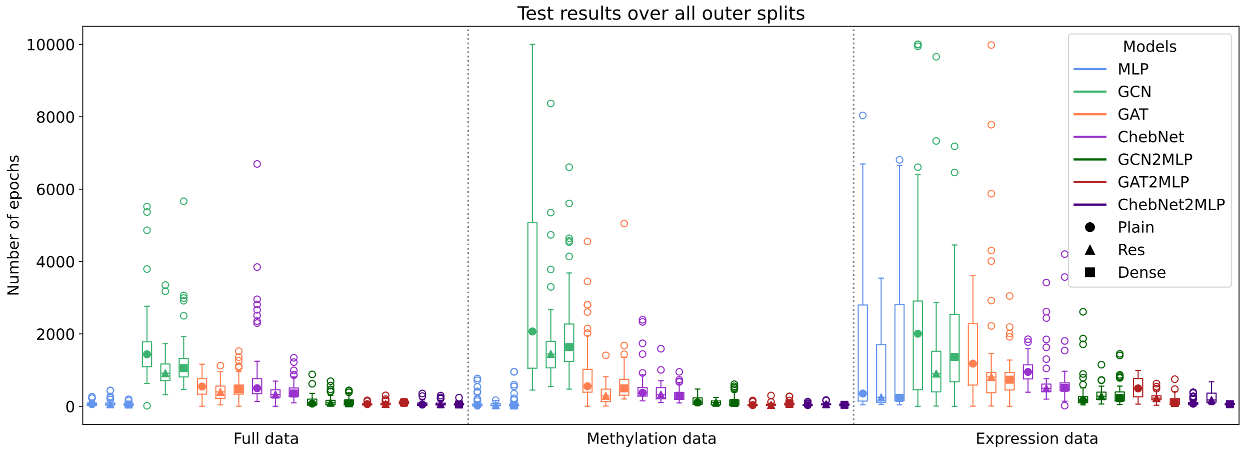


Figure 3: The distribution of training epoch numbers utilized per model architecture and data set for the **KIRC data**.

Table 1: Results per model architecture for the full data over 10 test sets in the form of the mean and the standard deviation of accuracy, AUROC, and F1 score for **KIRC data**.

| | Accuracy | AUROC | F1 score |
|---|---|---|---|
| MLP | | | |
| Plain | $90.10 \pm 1.99\%$ | $0.85 \pm 0.04$ | $0.92 \pm 0.01$ |
| Res | $89.47 \pm 2.24\%$ | $0.86 \pm 0.04$ | $0.92 \pm 0.01$ |
| Dense | $90.04 \pm 2.05\%$ | $0.86 \pm 0.04$ | $0.92 \pm 0.01$ |
| GCN | | | |
| Plain | $89.47 \pm 2.30\%$ | $0.86 \pm 0.03$ | $0.92 \pm 0.02$ |
| Res | $89.88 \pm 2.22\%$ | $0.87 \pm 0.04$ | $0.92 \pm 0.02$ |
| Dense | $89.88 \pm 2.08\%$ | $0.87 \pm 0.03$ | $0.92 \pm 0.01$ |
| GAT | | | |
| Plain | $85.92 \pm 4.44\%$ | $0.82 \pm 0.05$ | $0.90 \pm 0.03$ |
| Res | $87.90 \pm 6.18\%$ | $0.86 \pm 0.04$ | $0.91 \pm 0.04$ |
| Dense | $88.84 \pm 2.64\%$ | $0.86 \pm 0.03$ | $0.92 \pm 0.02$ |
| ChebNet | | | |
| Plain | $90.12 \pm 1.97\%$ | $0.88 \pm 0.03$ | $0.93 \pm 0.01$ |
| Res | $89.47 \pm 2.47\%$ | $0.87 \pm 0.03$ | $0.92 \pm 0.02$ |
| Dense | $89.47 \pm 2.48\%$ | $0.87 \pm 0.03$ | $0.92 \pm 0.02$ |
| GCN2MLP | | | |
| Plain | $90.10 \pm 1.99\%$ | $0.86 \pm 0.04$ | $0.92 \pm 0.01$ |
| Res | $90.10 \pm 1.99\%$ | $0.86 \pm 0.04$ | $0.92 \pm 0.01$ |
| Dense | $90.10 \pm 1.99\%$ | $0.86 \pm 0.04$ | $0.92 \pm 0.01$ |
| GAT2MLP | | | |
| Plain | $89.86 \pm 1.97\%$ | $0.87 \pm 0.03$ | $0.92 \pm 0.01$ |
| Res | $89.88 \pm 2.01\%$ | $0.87 \pm 0.04$ | $0.92 \pm 0.01$ |
| Dense | $87.08 \pm 3.39\%$ | $0.86 \pm 0.04$ | $0.90 \pm 0.03$ |
| ChebNet2MLP | | | |
| Plain | $90.10 \pm 1.99\%$ | $0.86 \pm 0.04$ | $0.92 \pm 0.01$ |
| Res | $89.92 \pm 2.23\%$ | $0.86 \pm 0.04$ | $0.92 \pm 0.02$ |
| Dense | $89.98 \pm 2.16\%$ | $0.86 \pm 0.04$ | $0.92 \pm 0.02$ |

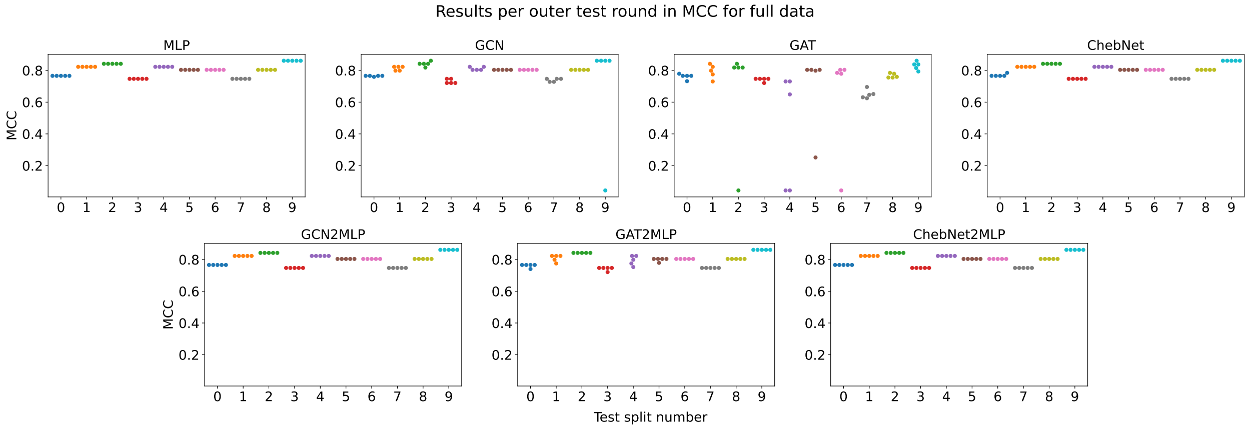


Figure 4: The exact MCC results per test split on the **KIRC data**. Every color represents results from one outer test split, whereas the 5 dots per color represent the 5 outer repetitions due to 5-fold CV.

#### 2.2 BRCA data

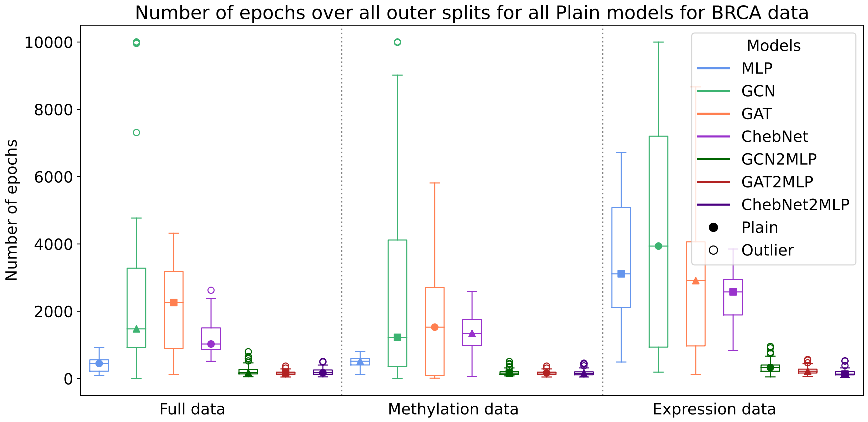


Figure 5: The distribution of training epoch numbers utilized per model architecture and data set for the **BRCA data**.

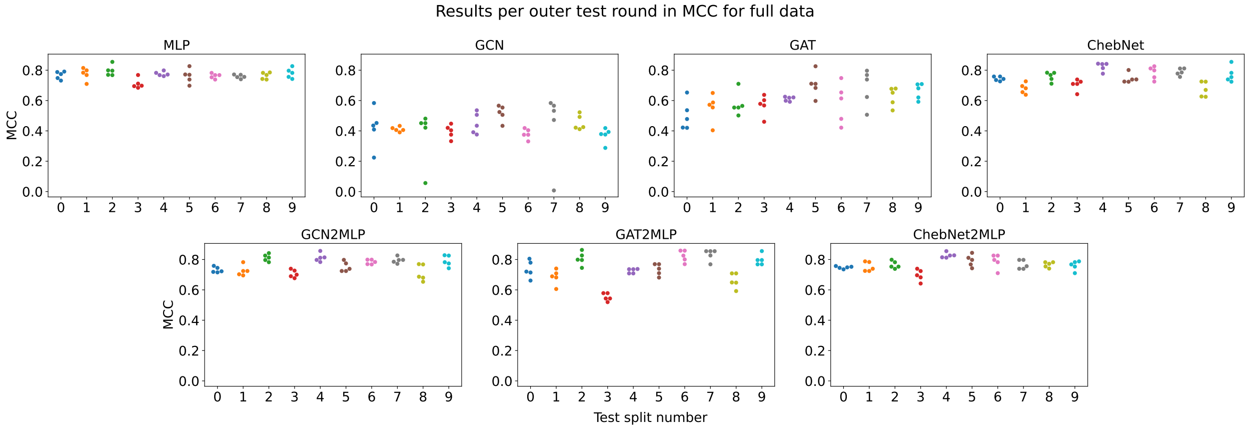


Figure 6: The exact MCC results per test split on the **BRCA data**. Every color represents results from one outer test split, whereas the 5 dots per color represent the 5 outer repetitions due to 5-fold CV.

Table 2: Results per model architecture for the full data over 10 test sets in the form of the mean and the standard deviation of accuracy, AUROC, and F1 score for **BRCA data**.

| | Accuracy | AUROC | F1 score |
|---|---|---|---|
| MLP | $88.17 \pm 1.12\%$ | $0.96 \pm 0.01$ | $0.89 \pm 0.01$ |
| GCN | $70.93 \pm 2.33\%$ | $0.78 \pm 0.02$ | $0.74 \pm 0.02$ |
| GAT | $80.26 \pm 3.27\%$ | $0.88 \pm 0.02$ | $0.81 \pm 0.04$ |
| ChebNet | $87.28 \pm 2.41\%$ | $0.95 \pm 0.01$ | $0.88 \pm 0.02$ |
| GCN2MLP | $88.00 \pm 2.13\%$ | $0.95 \pm 0.01$ | $0.89 \pm 0.02$ |
| GAT2MLP | $86.64 \pm 4.42\%$ | $0.94 \pm 0.02$ | $0.87 \pm 0.04$ |
| ChebNet2MLP | $87.28 \pm 2.41\%$ | $0.95 \pm 0.01$ | $0.88 \pm 0.02$ |

### 3 Details on hyperparameter selection

#### 3.1 KIRC data

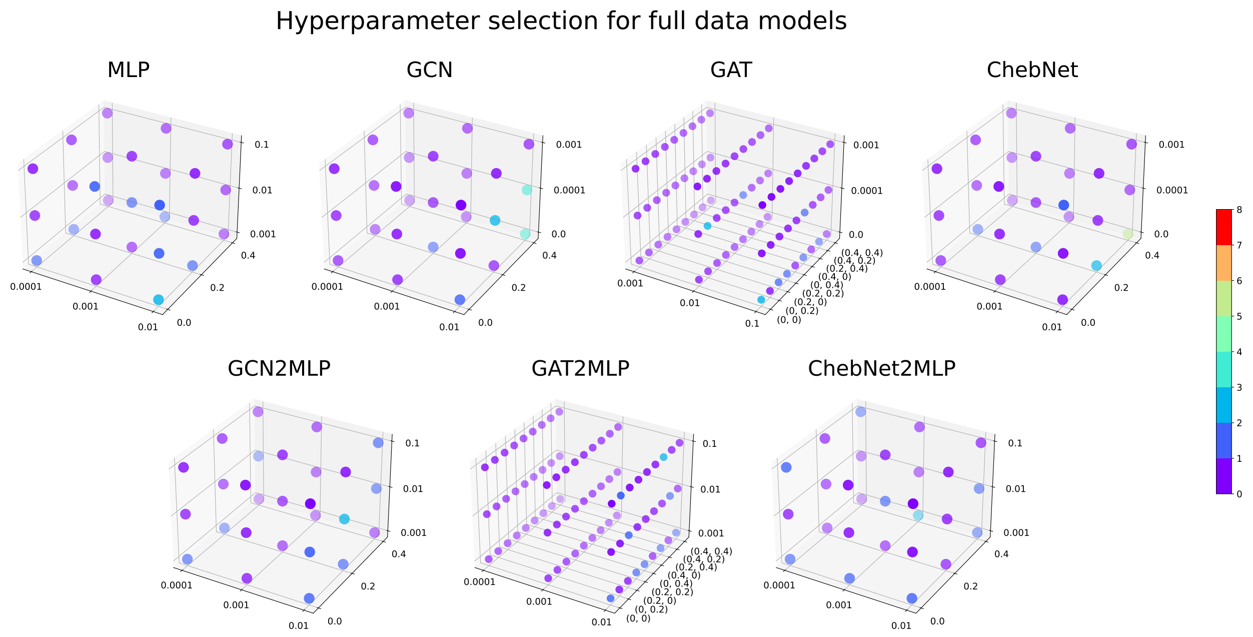


Figure 7: The number of selected hyperparameter combinations for the outer 10 test sets per model for Plain models on the full **KIRC data**.

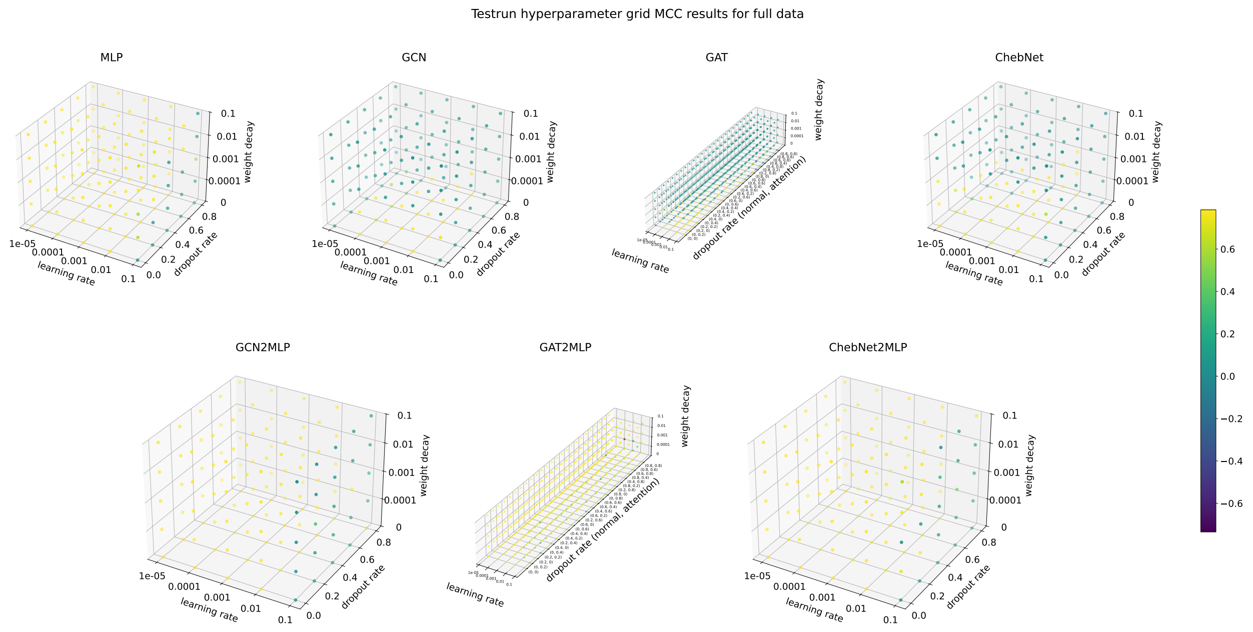


Figure 8: The MCC performance for the wide hyperparameter grid in the pretest for the plain models on the full **KIRC data**

Table 3: The hyperparameter ranges for the final benchmark on the **KIRC data**; note that the residual and dense variants of the models utilize the same hyperparameter ranges as the corresponding plain model. For the GAT models, the "both" in the dropout rate column indicates that the normal and attention head related dropout rates have the same possible choices.

| | learning rate | weight decay | dropout rate |
|---|---|---|---|
| MLP | 0.0001, 0.001, 0.01 | 0.001, 0.01, 0.1 | 0, 0.2, 0.4 |
| GCN | 0.0001, 0.001, 0.01 | 0, 0.0001, 0.001 | 0, 0.2, 0.4 |
| GAT | 0.001, 0.01, 0.1 | 0, 0.0001, 0.001 | 0, 0.2, 0.4 (both) |
| ChebNet | 0.0001, 0.001, 0.01 | 0, 0.0001, 0.001 | 0, 0.2, 0.4 |
| GCN2MLP | 0.0001, 0.001, 0.01 | 0.001, 0.01, 0.1 | 0, 0.2, 0.4 |
| GAT2MLP | 0.0001, 0.001, 0.01 | 0.001, 0.01, 0.1 | 0, 0.2, 0.4 (both) |
| ChebNet2MLP | 0.0001, 0.001, 0.01 | 0.001, 0.01, 0.1 | 0, 0.2, 0.4 |
| mMLP | 0.0001, 0.001, 0.01 | 0.001, 0.01, 0.1 | 0, 0.2, 0.4 |
| mGCN | 0.0001, 0.001, 0.01 | 0, 0.0001, 0.001 | 0, 0.2, 0.4 |
| mGAT | 0.001, 0.01, 0.1 | 0, 0.0001, 0.001 | 0, 0.2, 0.4 (both) |
| mChebNet | 0.0001, 0.001, 0.01 | 0, 0.0001, 0.001 | 0, 0.2, 0.4 |
| mGCN2MLP | 0.0001, 0.001, 0.01 | 0.001, 0.01, 0.1 | 0, 0.2, 0.4 |
| mGAT2MLP | 0.0001, 0.001, 0.01 | 0.001, 0.01, 0.1 | 0, 0.2, 0.4 (both) |
| mChebNet2MLP | 0.0001, 0.001, 0.01 | 0.0001, 0.001, 0.01 | 0, 0.2, 0.4 |
| xMLP | 0.00001, 0.0001, 0.001 | 0.001, 0.01, 0.1 | 0, 0.2, 0.4 |
| xGCN | 0.0001, 0.001 , 0.01 | 0, 0.0001, 0.001 | 0, 0.2, 0.4 |
| xGAT | 0.0001, 0.001, 0.01 | 0, 0.0001, 0.001 | 0, 0.2, 0.4 (both) |
| xChebNet | 0.0001, 0.001, 0.01 | 0, 0.0001, 0.001 | 0, 0.2, 0.4 |
| xGCN2MLP | 0.0001, 0.001, 0.01 | 0.0001, 0.001, 0.01 | 0, 0.2, 0.4 |
| xGAT2MLP | 0.0001, 0.001, 0.01 | 0.001, 0.01, 0.1 | 0, 0.2, 0.4 (both) |
| xChebNet2MLP | 0.0001, 0.001, 0.01 | 0.0001, 0.001, 0.01 | 0, 0.2, 0.4 |

#### 3.2 BRCA data

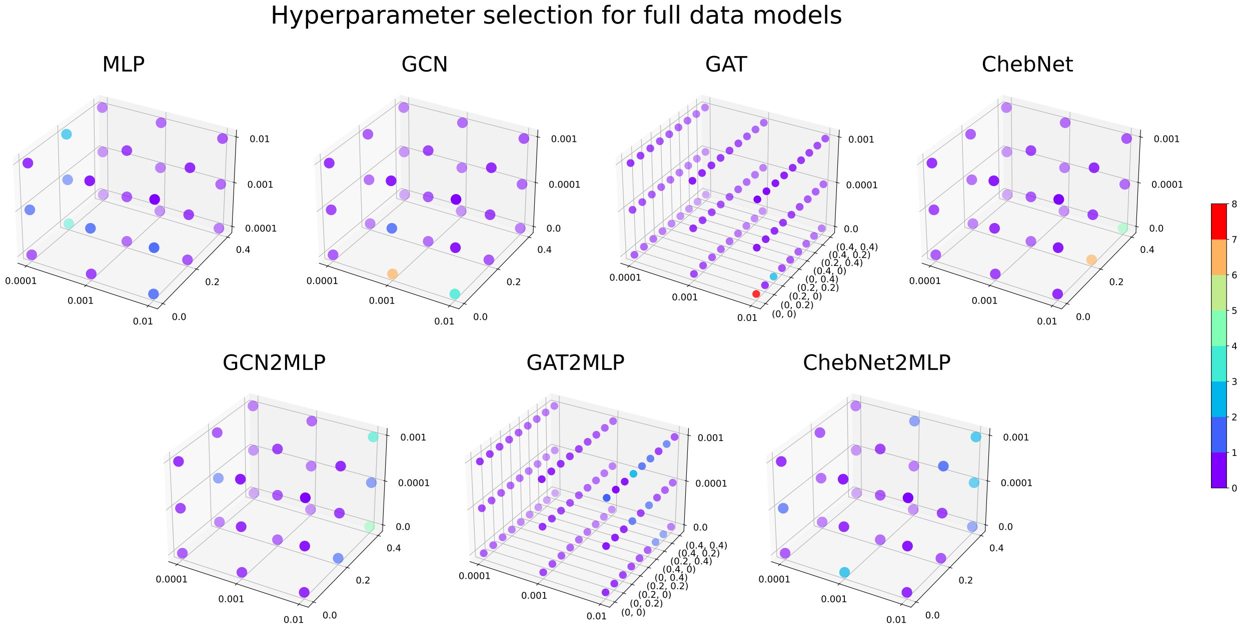


Figure 9: The number of selected hyperparameter combinations for the outer 10 test sets per model for Plain models on the full **BRCA data**.

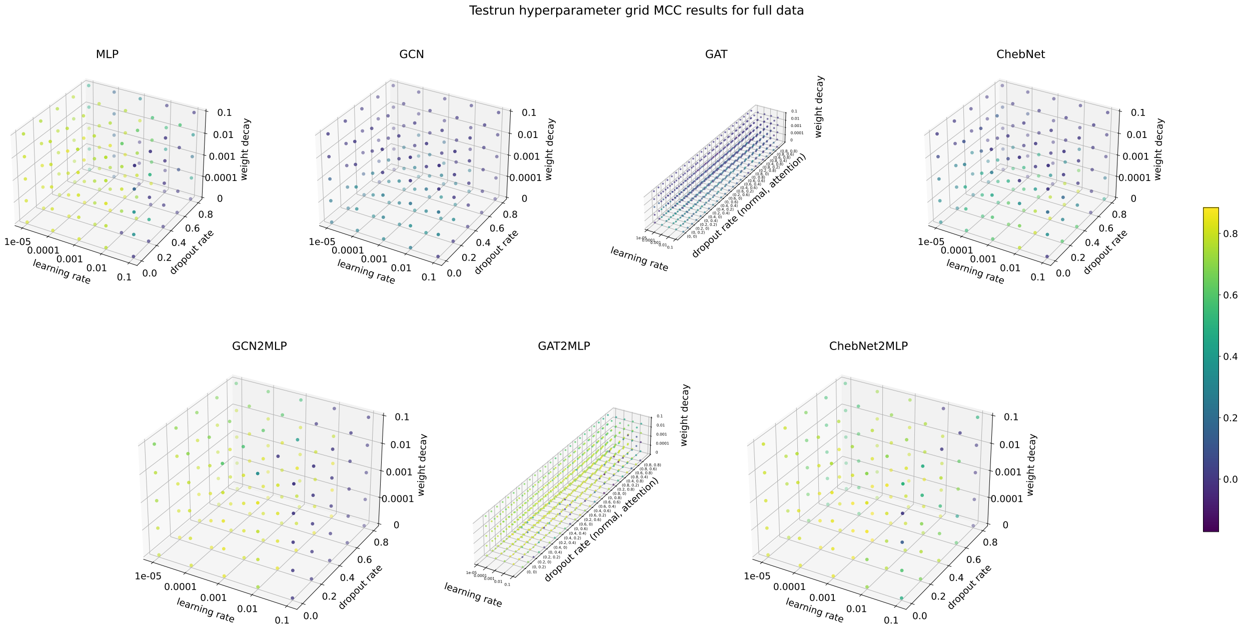


Figure 10: The MCC performance for the wide hyperparameter grid in the pretest for the plain models on the full **BRCA data**

Table 4: The hyperparameter ranges for the final benchmark on the **BRCA data**; note that the residual and dense variants of the models utilize the same hyperparameter ranges as the corresponding plain model. For the GAT models, the "both" in the dropout rate column indicates that the normal and attention head related dropout rates have the same possible choices.

| | learning rate | weight decay | dropout rate |
|---|---|---|---|
| MLP | 0.0001, 0.001, 0.01 | 0.0001, 0.001, 0.01 | 0, 0.2, 0.4 |
| GCN | 0.0001, 0.001, 0.01 | 0, 0.0001, 0.001 | 0, 0.2, 0.4 |
| GAT | 0.0001, 0.001, 0.01 | 0, 0.0001, 0.001 | 0, 0.2, 0.4 (both) |
| ChebNet | 0.0001, 0.001, 0.01 | 0, 0.0001, 0.001 | 0, 0.2, 0.4 |
| GCN2MLP | 0.0001, 0.001, 0.01 | 0, 0.0001, 0.001 | 0, 0.2, 0.4 |
| GAT2MLP | 0.0001, 0.001, 0.01 | 0, 0.0001, 0.001 | 0, 0.2, 0.4 (both) |
| ChebNet2MLP | 0.0001, 0.001, 0.01 | 0, 0.0001, 0.001 | 0, 0.2, 0.4 |
| mMLP | 0.0001, 0.001, 0.01 | 0.0001, 0.001, 0.01 | 0, 0.2, 0.4 |
| mGCN | 0.0001, 0.001, 0.01 | 0, 0.0001, 0.001 | 0, 0.2, 0.4 |
| mGAT | 0.0001, 0.001, 0.01 | 0, 0.0001, 0.001 | 0, 0.2, 0.4 (both) |
| mChebNet | 0.001, 0.01, 0.1 | 0, 0.0001, 0.001 | 0, 0.2, 0.4 |
| mGCN2MLP | 0.0001, 0.001, 0.01 | 0, 0.0001, 0.001 | 0, 0.2, 0.4 |
| mGAT2MLP | 0.0001, 0.001, 0.01 | 0, 0.0001, 0.001 | 0, 0.2, 0.4 (both) |
| mChebNet2MLP | 0.0001, 0.001, 0.01 | 0, 0.0001, 0.001 | 0, 0.2, 0.4 |
| xMLP | 0.00001, 0.0001, 0.001 | 0.0001, 0.001, 0.01 | 0, 0.2, 0.4 |
| xGCN | 0.0001, 0.001 , 0.01 | 0, 0.0001, 0.001 | 0, 0.2, 0.4 |
| xGAT | 0.0001, 0.001, 0.01 | 0, 0.0001, 0.001 | 0, 0.2, 0.4 (both) |
| xChebNet | 0.001, 0.01, 0.1 | 0, 0.0001, 0.001 | 0, 0.2, 0.4 |
| xGCN2MLP | 0.0001, 0.001, 0.01 | 0, 0.0001, 0.001 | 0, 0.2, 0.4 |
| xGAT2MLP | 0.0001, 0.001, 0.01 | 0, 0.0001, 0.001 | 0, 0.2, 0.4 (both) |
| xChebNet2MLP | 0.0001, 0.001, 0.01 | 0, 0.0001, 0.001 | 0, 0.2, 0.4 |

### 4 Number of model parameters

Table 5: Number of model parameters, i.e., the weights and biases, per model exemplarily for the **KIRC data**.

| | methylation/expression data | full data |
|---|---|---|
| MLP | | |
| Plain | 106 305 | 208 321 |
| Res | 110 465 | 212 481 |
| Dense | 107 393 | 209 409 |
| GCN | | |
| Plain | 4 353 | 4 417 |
| Res | 8 513 | 8 576 |
| Dense | 5 441 | 5 505 |
| GAT | | |
| Plain | 4 609 | 4 673 |
| Res | 8 897 | 8 961 |
| Dense | 5 697 | 5 761 |
| ChebNet | | |
| Plain | 33 473 | 33 985 |
| Res | 66 305 | 66 817 |
| Dense | 41 729 | 42 241 |
| GCN2MLP | | |
| Plain | 106 305 | 106 369 |
| Res | 110 465 | 110 528 |
| Dense | 209 345 | 209 409 |
| GAT2MLP | | |
| Plain | 106 561 | 106 625 |
| Res | 110 849 | 110 913 |
| Dense | 209 601 | 209 665 |
| ChebNet2MLP | | |
| Plain | 135 425 | 135 937 |
| Res | 168 257 | 168 769 |
| Dense | 245 663 | 246 145 |
